# HB-PLS: An algorithm for identifying biological process or pathway regulators by integrating Huber loss and Berhu penalty with partial least squares regression

**DOI:** 10.1101/2020.05.16.089623

**Authors:** Wenping Deng, Kui Zhang, Zhigang Wei, Lihu Wang, Cheng He, Sanzhen Liu, Hairong Wei

## Abstract

Gene expression data features high dimensionality, multicollinearity, and the existence of outlier or non-Gaussian distribution noise, which make the identification of true regulatory genes controlling a biological process or pathway difficult. In this study, we embedded the Huber-Berhu (HB) regression into the partial least squares (PLS) framework and created a new method called HB-PLS for predicting biological process or pathway regulators through construction of regulatory networks. PLS is an alternative to ordinary least squares (OLS) for handling multicollinearity in high dimensional data. The Huber loss is more robust to outliers than square loss, and the Berhu penalty can obtain a better balance between the *ℓ*_2_ penalty and the *ℓ*_1_ penalty. HB-PLS therefore inherits the advantages of the Huber loss, the Berhu penalty, and PLS. To solve the Huber-Berhu regression, a fast proximal gradient descent method was developed; the HB regression runs much faster than CVX, a Matlab-based modeling system for convex optimization. Implementation of HB-PLS to real transcriptomic data from *Arabidopsis* and maize led to the identification of many pathway regulators that had previously been identified experimentally. In terms of its efficiency in identifying positive biological process or pathway regulators, HB-PLS is comparable to sparse partial least squares (SPLS), a very efficient method developed for variable selection and dimension reduction in handling multicollinearity in high dimensional genomic data. However, HB-PLS is able to identify some distinct regulators, and in one case identify more positive regulators at the top of output list, which can reduce the burden for experimental test of the identified candidate targets. Our study suggests that HB-PLS is instrumental for identifying biological process and pathway genes.

## Introduction

In a gene regulatory network (GRN), a node corresponds to a gene and an edge represents a directional regulatory relationship between a transcription factor (TF) and a target gene. Understanding the regulatory relationships among genes in GRNs can help elucidate the various biological processes and underlying mechanisms in a variety of organisms. Although experiments can be conducted to acquire evidence of gene regulatory interactions, these are labor-intensive and time-consuming. In the past two decades, the advent of high-throughput techniques, including microarray and RNA-Seq, have generated an enormous wealth of transcriptomic data. As the data in public repositories grows exponentially, computational algorithms and tools utilizing gene expression data offer a more time- and cost-effective way to reconstruct GRNs. To this end, efficient mathematical and statistical methods are needed to infer qualitative and quantitative relationships between genes.

Many methods have been developed to reconstruct GRNs, each employing different theories and principles. The earliest methods involved differential equations [1], Boolean networks [2], stochastic networks [3], Bayesian [4, 5] or dynamic Bayesian networks (BN) [6, 7], and ordinary differential equations (ODE) [8]. Some of these methods require time series datasets with short time intervals, such as those generated from easily manipulated single cell organisms (e.g. bacteria, yeast) or mammalian cell lines [9]. For this reason, most of these methods are not suitable for gene expression data, especially time series data involving time intervals on the scale of days, from multicellular organisms like plants and mammals (except cell lines).

In general, the methods that are useful for building gene networks with non-time series data generated from high plants and mammals include ParCorA [10], GGM [11], and mutual information-based methods such as Relevance Network (RN) [12], Algorithm for the Reconstruction of Accurate Cellular Networks (ARACNE) [13], C3NET [14], maximum relevance/minimum redundancy Network (MRNET) [15], and random forests [16, 17]. Most of these methods use the information-theoretic framework. For instance, Relevance Network (RN) [18], one of the earliest methods, infers a network in which a pair of genes are linked by an edge if the mutual information is larger than a given threshold. The context likelihood relatedness (CLR) algorithm [19], an extension of RN, derives a score from the empirical distribution of the mutual information for each pair of genes and eliminates edges with scores that are not statistically significant. ARACNE [20] is similar to RN; however, ARACNE makes use of the data processing inequality (DPI) to eliminate the least significant edge of a triplet of genes, which decreases the false positive rate of the inferred network. MRNET [21] employs the maximum relevance and minimum redundancy feature selection method to infer GRNs. Finally, triple-gene mutual interaction (TGMI) uses condition mutual information to evaluate triple gene blocks to infer gene regulatory networks [22]. Information theory-based methods are used extensively for constructing GRNs and for building large networks because they have a low computational complexity and are able to capture nonlinear dependencies. However, there are also disadvantages in using mutual information, including high false-positive rates [23] and the inability to differentiate positive (activating), negative (inhibiting), and indirect regulatory relationships. Reconstruction of the transcriptional regulatory network can be implemented by the neighborhood selection method. Neighborhood selection [24] is a sub-problem of covariance selection. Assume Γ is a set containing all of the variables (genes), the neighborhood *ne*_*a*_ of a variable *a* ∈ Γ is the smallest subset of Γ\{*a*} such that, given all variables in *ne*_*a*_, variable *a* is conditionally independent of all remaining variables. Given *n* i.i.d. observations of Γ, neighborhood selection aims to estimate the neighborhood of each variable in Γ individually. The neighborhood selection problem can be cast as a multiple linear regression problem and solved by regularized methods.

Following the differential equation in [25], the expression levels of a target gene *y* and the expression levels of the TF genes *x* form a linear relationship:

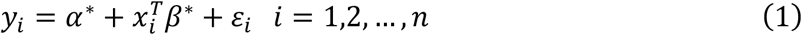

where *n* is the number of samples, *x*_*i*_ = (*x*_*i*1_, … , *x*_*ip*_)^*T*^ is the expression level of *p* TF genes, and *y*_*i*_ is the expression level of the target gene in sample *i*. *α*\* is the intercept and 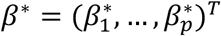 are the associated regression coefficients; if 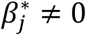, then TF gene *j* regulates target gene *i*. {*ε*_*i*_} are independent and identically distributed random errors with mean 0 and variance *σ*^2^. The method to get an approximation 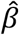 for *β*\* is to transform this statistical problem to a convex optimization problem:

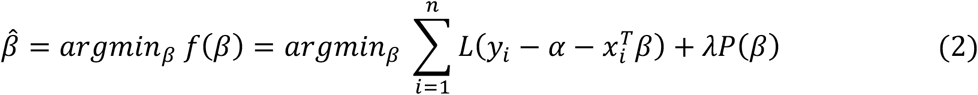

where *L*(·) is a loss function, *P*(·) is a penalization function, and *λ* > 0 is a tuning parameter which determines the importance of penalization. Different loss functions, penalization functions, and methods for determining *λ* have been proposed in the literature. Ordinary least squares (OLS) is the simplest method with a square loss function 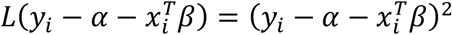 and no penalization function. The OLS estimator is unbiased. However, since it is common for the number of genes, *p*, *to* be much larger than the number of samples, *n*, (i.e. *p* ≫ *n*) in any given gene expression data set, there is no unique solution for OLS. Even when *n* > *p*, OLS estimation features high variance. To conquer these problems, ridge regression [26] adds a *ℓ*2 penalty, 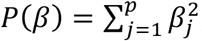, on the coefficients which introduces a bias but reduces the variance of the estimated 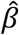. In ridge regression, there is a unique solution even for the *p* > *n* case. Least absolute shrinkage and selection operator [27] is similar to ridge regression, except the *ℓ*2 penalty in ridge regression is replaced by the *ℓ*1 penalty, 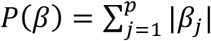.

The main benefit of LASSO is that it performs variable selection and regularization simultaneously thereby generating a sparse solution, a desirable property for constructing GRNs. When LASSO is used for selecting regulatory TFs for a target gene, there are two potential limitations. First, if several TF genes are correlated and have large effects on the target gene, LASSO has a tendency to choose only one TF gene while zeroing out the other TF genes. Second, some studies [28] state that LASSO does not have oracle properties; that is, it does not have the capability to identify the correct subset of true variables or to have an optimal estimation rate. It is claimed that there are cases where a given *λ* that leads to optimal estimation rate ends up with an inconsistent selection of variables. For the first limitation, Zou and Hastie [29] proposed elastic net, in which the penalty is a mixture of LASSO and ridge regressions: 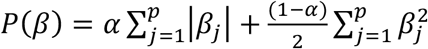, where *α* (0 < *α* < 1) is called the elastic net mixing parameter. When *α* = 1, the elastic net penalty becomes the LASSO penalty; when *α* = 0, the elastic net penalty becomes the ridge penalty. For the second limitation, adaptive LASSO [28] was proposed as a regularization method, which enjoys the oracle properties. The penalty function for adaptive LASSO is: 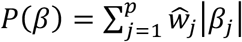, where adaptive weight 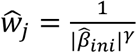, and 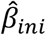 is an initial estimate of the coefficients obtained through ridge regression or LASSO; *γ* is a positive constant, and is usually set to 1. It is evident that adaptive LASSO penalizes more those coefficients with lower initial estimates.

It is well known that the square loss function is sensitive to heavy-tailed errors or outliers. Therefore, adaptive LASSO may fail to produce reliable estimates for datasets with heavy-tailed errors or outliers, which commonly appear in gene expression datasets. One possible remedy is to remove influential observations from the data before fitting a model, but it is difficult to differentiate true outliers from normal data. The other method is to use robust regression. Wang et al. [30] combined the least absolute deviation (LAD) and weighted LASSO penalty to produce the LAD-LASSO method. The objective function is:

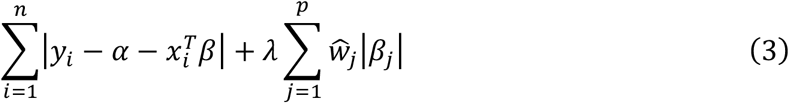

With this LAD loss, LAD-LASSO is more robust than OLS to unusual *y* values, but it is sensitive to high leverage outliers. Moreover, LAD estimation degrades the efficiency of the resulting estimation if the error distribution is not heavy tailed [31]. To achieve both robustness and efficiency, Lambert-Lacroix et al. [32] proposed Huber-LASSO, which combined the Huber loss function and a weighted LASSO penalty. The Huber function (see Materials and Methods) is a hybrid of squared error for relatively small errors and absolute error for relatively large ones. Owen [33] proposed the use of the Huber function as a loss function and the use of a reversed version of Huber’s criterion, called Berhu, as a penalty function. For the Berhu penalty (see Materials and Methods), relatively small coefficients contribute their *ℓ*1 norm to the penalty while larger ones cause it to grow quadratically. This Berhu penalty sets some coefficients to 0, like LASSO, while shrinking larger coefficients in the same way as ridge regression. In [34], the authors showed that the combination of the Huber loss function and an adaptive Berhu penalty enjoys oracle properties, and they also demonstrated that this procedure encourages a grouping effect. In [33, 34], the authors solved a Huber-Berhu optimization problem using CVX software [35], a Matlab-based modeling system for convex optimization. CVX turns Matlab into a modeling language, allowing constraints and objectives to be specified using standard Matlab expression syntax. However, since CVX is slow for large datasets, a proximal gradient descent algorithm was developed for the Huber-Berhu regression in this study, which runs much faster than CVX.

Reconstruction of gene regulatory networks often involves ill-posed problems due to high dimensionality and multicollinearity. Partial least squares (PLS) regression has been an alternative to ordinary regression for handling multicollinearity in several areas of scientific research. PLS couples a dimension reduction technique and a regression model. Although PLS has been shown to have good predictive performance in dealing with ill-posed problems, it is not particularly tailored for variable selection. Chun et al. [36] first proposed a SPLS regression for simultaneous dimension reduction and variable selection. Cao et al. [37] also proposed a sparse PLS method for variable selection when integrating omics data. They added sparsity into PLS with a LASSO penalization combined with singular value decomposition (SVD) computation. In this study, the Huber-Berhu regression was embedded into a PLS framework. Real gene data was used to demonstrate that this approach is applicable for the reconstruction of GRNs.

## Materials and Methods

### High-throughput gene expression data

The lignin pathway analysis used an *Arabidopsis* wood formation compendium dataset containing 128 Affymetrix microarrays pooled from six experiments (accession identifiers: GSE607, GSE6153, GSE18985, GSE2000, GSE24781, and GSE5633 in NCBI Gene Expression Omnibus (GEO) (http://www.ncbi.nlm.nih.gov/geo/)). These datasets were originally obtained from hypocotyledonous stems under short-day conditions known to induce secondary wood formation [38]. The original CEL files were downloaded from GEO and preprocessed using the affy package in Bioconductor (https://www.bioconductor.org/) and then normalized with the robust multi-array analysis (RMA) algorithm in affy package. This compendium data set was also used in our previous studies [39, 40]. The maize B73 compendium data set used for predicting photosynthesis light reaction (PLR) pathway regulators was downloaded from three NCBI databases: (1) the sequence read archive (SRA) (https://www.ncbi.nlm.nih.gov/sra), 39 leaf samples from ERP011838; (2) Gene Expression Omnibus (GEO), 24 leaf samples from GSE61333, and (3) BioProject (https://www.ncbi.nlm.nih.gov/bioproject/), 36 seedling samples from PRJNA483231. This compendium is a subset of that used in our earlier co-expression analysis [41]. Raw reads were trimmed to remove adaptors and low-quality base pairs via Trimmomatic (v3.3). Clean reads were aligned to the B73Ref3 with STAR, followed by the generation of normalized FPKM (fragments per kb of transcript per million reads) using Cufflinks software (v2.1.1) [42].

### Huber and Berhu functions

In estimating regression coefficients, the square loss function is well suited if *y*_*i*_ follows a Gaussian distribution, but it gives a poor performance when *y*_*i*_ follows a heavy-tailed distribution or there are outliers. On the other hand, the LAD loss function is more robust to outliers, but the statistical efficiency is low when there are no outliers in the data. The Huber function, introduced in [43], is a combination of linear and quadratic loss functions. For any given positive real *M* (called shape parameter), the Huber function is defined as:

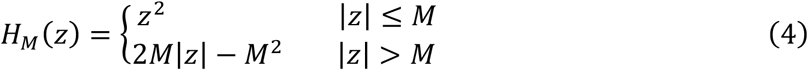

This function is quadratic for small *z* but grows linearly for large values of *z*. The parameter *M* determines where the transition from quadratic to linear takes place (see Figure 1, top left). In this study, the default value of *M* was set to be one tenth of the interquartile range (IRQ), as suggested by [44]. The Huber function is a smooth function with a derivative function:

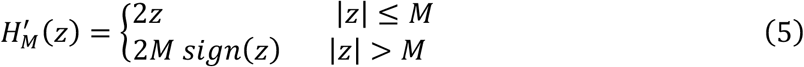

**Figure 1.**
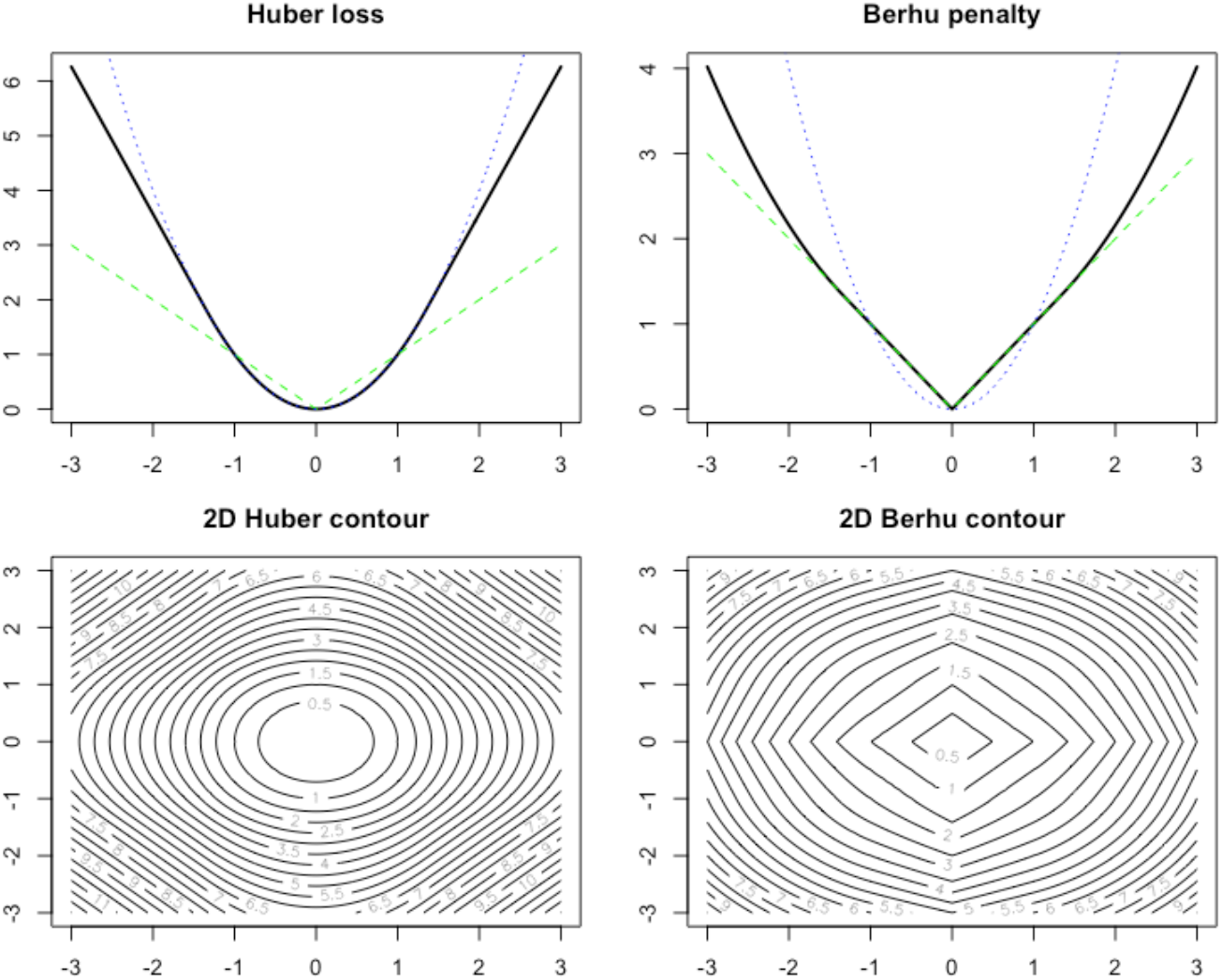
Huber loss function (top left) and Berhu penalty function (top right) as well as their 2D contours (bottom row).

The ridge regression uses the quadratic penalty on regression coefficients, and it is equivalent to putting a Gaussian prior on the coefficients. LASSO uses a linear penalty on regression coefficients, and it is equivalent to putting a Laplace prior on the coefficients. The advantage of LASSO over ridge regression is that it implements regularization and variable selection simultaneously. The disadvantage is that, if a group of predictors is highly correlated, LASSO picks only one of them and shrinks the others to zero. In this case, the prediction performance of ridge regression dominates the LASSO. The Berhu function, introduced in [33], is a hybrid of these two penalties. It gives a quadratic penalty to large coefficients while giving a linear penalty to small coefficients, as shown in Figure 1 (top right). The Berhu function is defined as:

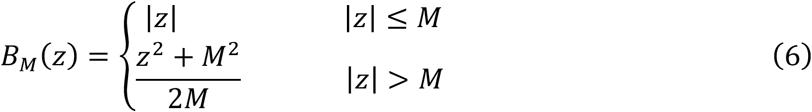

The shape parameter *M* was set to be the same as that in the Huber function. As shown in Figure 1 (top right), the Berhu function is a convex function, but it is not differentiable at *z* = 0. Figure 1 (bottom) also shows the 2D contour of Huber and Berhu functions. When the Huber loss function and the Berhu penalty were combined, an objective function, as referred as Huber_Berhu, was obtained, as shown below.

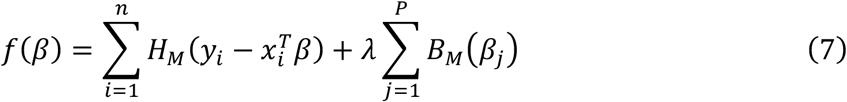

Figure 2 provides insight into the estimation of coefficients for the Huber_Berhu (left), LASSO (middle), and ridge (right) regressions. The Huber loss corresponds to the rotated, rounded rectangle contour in the top right corner, and the center of the contour is the solution of the unpenalized Huber regression. The shaded area is a map of the Berhu constraint where a smaller *λ* corresponds to a larger area. The estimated coefficient of the Huber_Berhu regression is the first place the contours touch the shaded area; when *λ* is small, the touch point is not on the axes, which means the Huber_Berhu regression behaves more like the ridge regression, which does not generate a sparse solution. When *λ* increases, the correspondent shaded area changes to a diamond, and the touch point is more likely to be located on the axes. Therefore, for large *λ*, the Huber_Berhu regression behaves like Lasso, which can generate a sparse solution.

**Figure 2.**
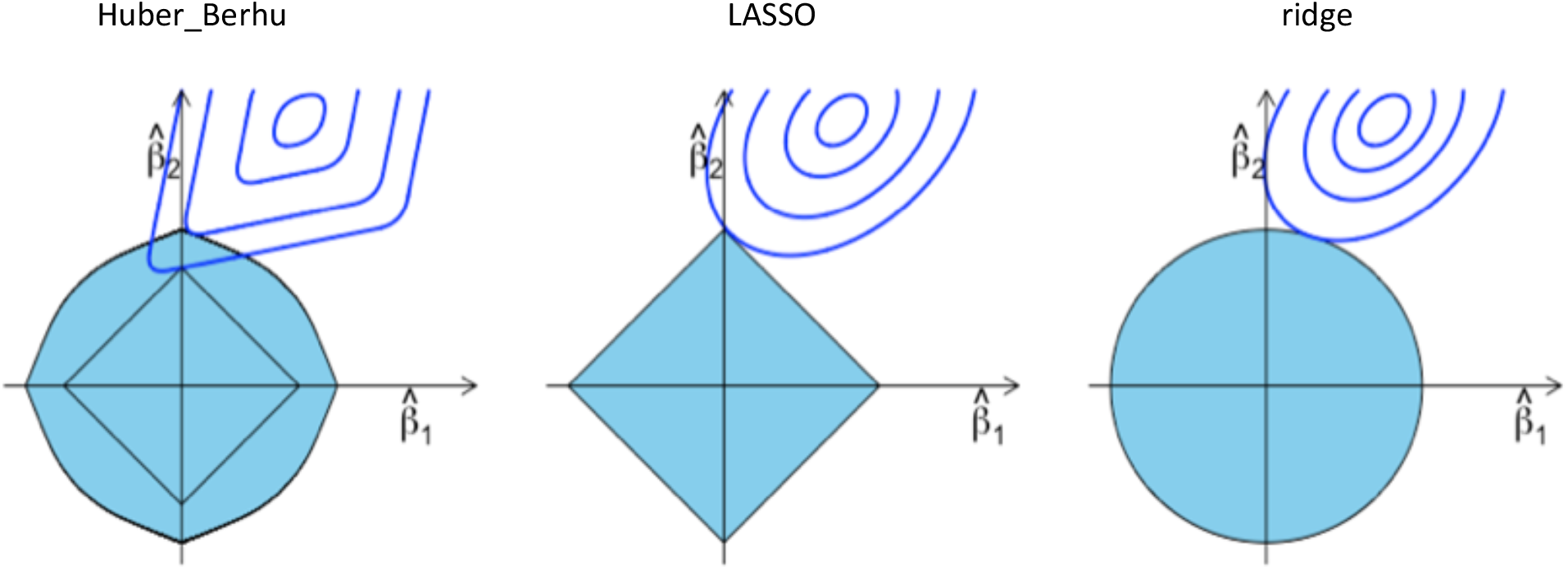
Estimation picture for the Huber_Berhu regression (left). As a comparison, the estimation pictures for the LASSO (middle) and ridge (right) regressions are also shown.

### The algorithm to solve the Huber-Berhu regression

Since the Berhu function is not differentiable at *z* = 0, it is difficult to use the gradient descent method to solve equation (4). Although we can use the general convex optimization solver CVX [35] for a convex optimization problem, it is too slow for real biological applications. Therefore, a proximal gradient descent algorithm was developed to solve equation (4). Proximal gradient descent is an effective algorithm to solve an optimization problem with decomposable objective function. Suppose the objective function can be decomposed as *f*(*x*) = *g*(*x*) + *h*(*x*), where *g*(*x*) is a convex differentiable function and *h*(*x*) is a convex non-differentiable function. The idea behind the proximal gradient descent [45] method is to make a quadratic approximation to *g*(*x*) and leave *h*(*x*) unchanged. That is:

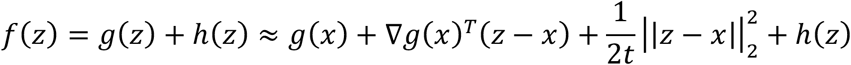

At each step, *x* is updated by the minimum of the right side of (5).

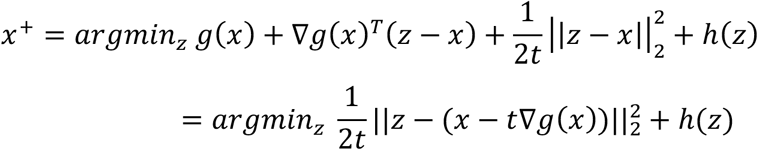

The operator 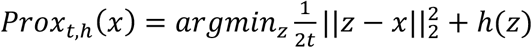 is called proximal mapping for *h*. Therefore to solve (4), the key is to compute the proximal mapping for the Berhu function:

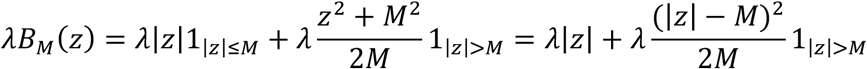

let 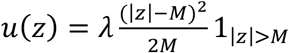. As *u*(*z*) satisfies theorem 4 in [46]:

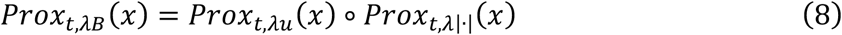

It is not difficult to verify:

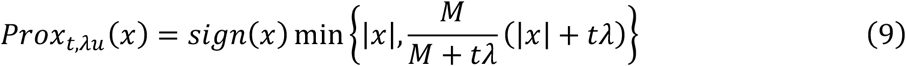

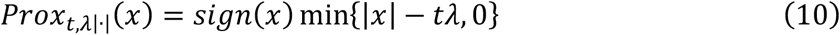

**Algorithm 1:**
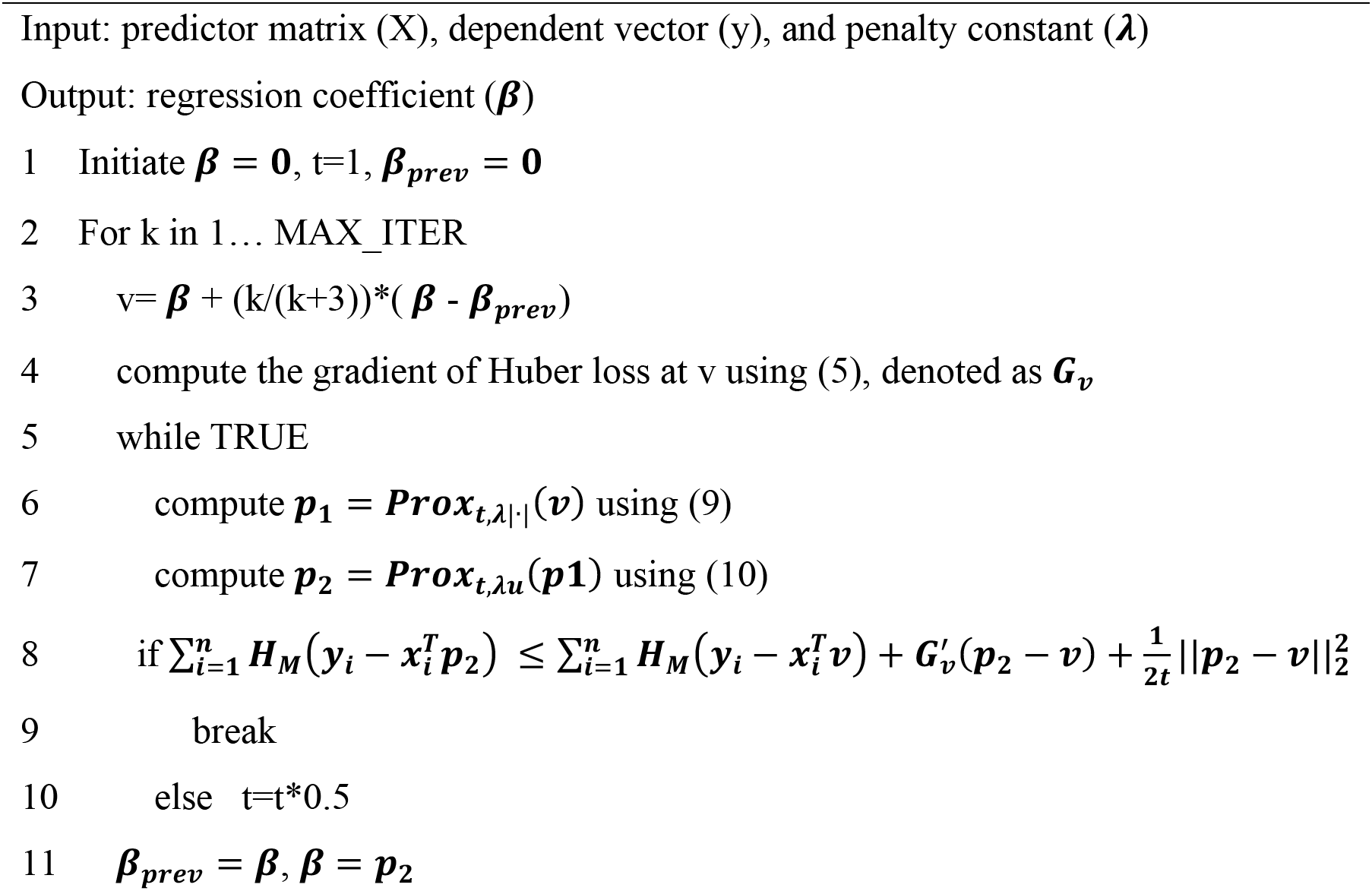
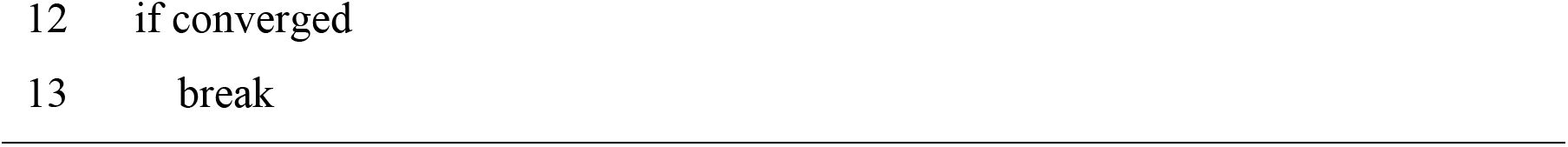
Accelerated proximal gradient descent method to solve equation (7)

Algorithm 1 uses the accelerated proximal gradient descent method to solve (7). Line 3 implements the acceleration of [47]. Lines 6-7 compute the proximal mapping of the Berhu function. Lines 5-10 use a backtracking method to determine the step size.

### Embedding the Huber-Berhu regression into PLS

Let *X* (*n* × *p*) and *Y* (*n* × *q*) be the standardized predictor variables (TF genes) and dependent variables (pathway genes), respectively. PLS [48] looks for a linear combination of *X* and a linear combination of *Y* such that their covariance reaches a maximum:

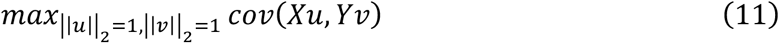

Here, the linear combination *ξ* = *Xu* and *η* = *Yv* are called component scores (or latent variables) and the *p* and *q* dimensional combinatory coefficients *u* and *v* are called loadings. After getting this first component *ξ*, two regression equations (from *X* to *ξ* and from *Y* to *ξ*) were set up:

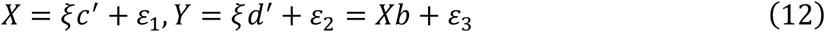

Next, X was deflated as X = X − ξc′ and Y was deflated as Y = Y − ξd′, and this process was continued until enough components were extracted.

A close relationship exists between PLS and SVD. Let *M* = *X*′*Y*, then 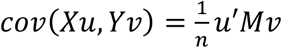. Let the SVD of *M* be:

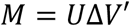

where *U*(*p* × *r*) and *V*(*q* × *r*) are orthonormal and Δ(*r* × *r*) is a diagonal matrix whose diagonal elements *δ*_*k*_ (*k* = 1 … *r*) are called singular values. According to the property of SVD, the combinatory coefficients *u* and *v* in (7) are exactly the first column of *U* and the first column of *V*. Therefore, the loadings of PLS can be computed by:

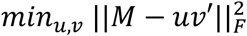

where 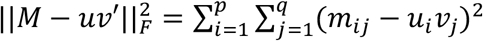.

Cao et al. [37] proposed a sparse PLS approach using SVD decomposition of *M* by adding a *ℓ*_1_ penalty on the loadings. The optimization problem to solve is:

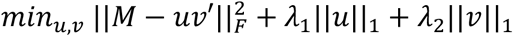

As mentioned above, the Huber function is more robust to outliers and has higher statistical efficiency than LAD loss, and the Berhu penalty has a better balance between the *ℓ*_1_ and *ℓ*_2_ penalty. The Huber loss and the Berhu penalty were adopted to extract each component for PLS. The optimization problem becomes:

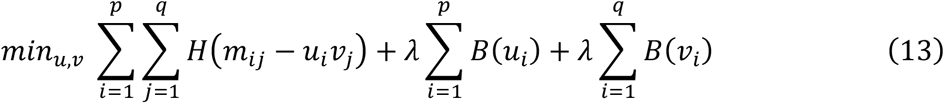

The objective function in (13) is not convex on *u* and *v*, but it is convex on *u* when *v* is fixed and convex on *v* when *u* is fixed. For example, when *v* is fixed, each *u*_*i*_ in parallel can be solved by:

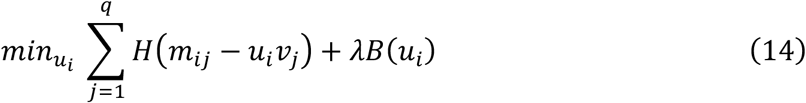

Similarly, when *u* is fixed, each *v*_*j*_ in parallel can be computed by:

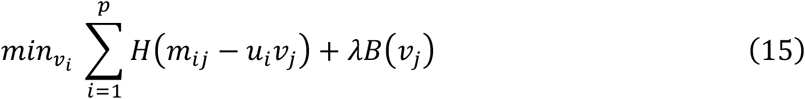

Equations (14) and (15) can be solved using Algorithm 1. Therefore (9) can be solved iteratively by updating *u* and *v* alternately. Note, it is not cost-efficient to spend a lot of effort optimizing over *u* in line 6 before a good estimate for *v* is computed. Since Algorithm 1 is an iterative algorithm, it may make sense to stop the optimization over *u* early before updating *v*. In the implementation, one step of proximal mapping was used to update *u* and *v*. That is:

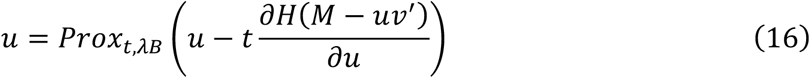

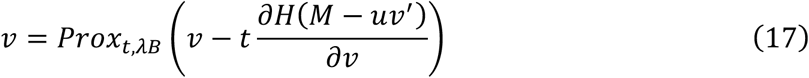

The Huber–Berhu PLS regression is detailed in Algorithm 2.

**Algorithm 2:**
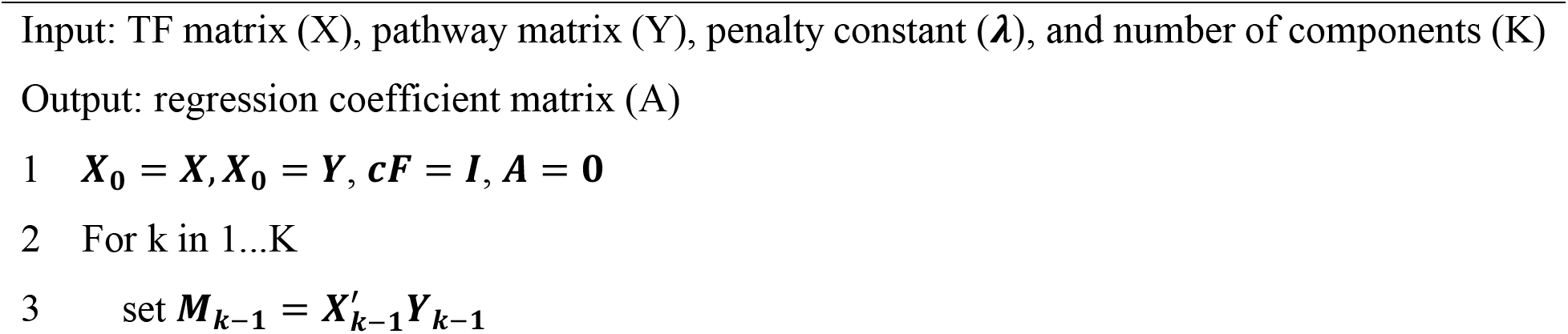
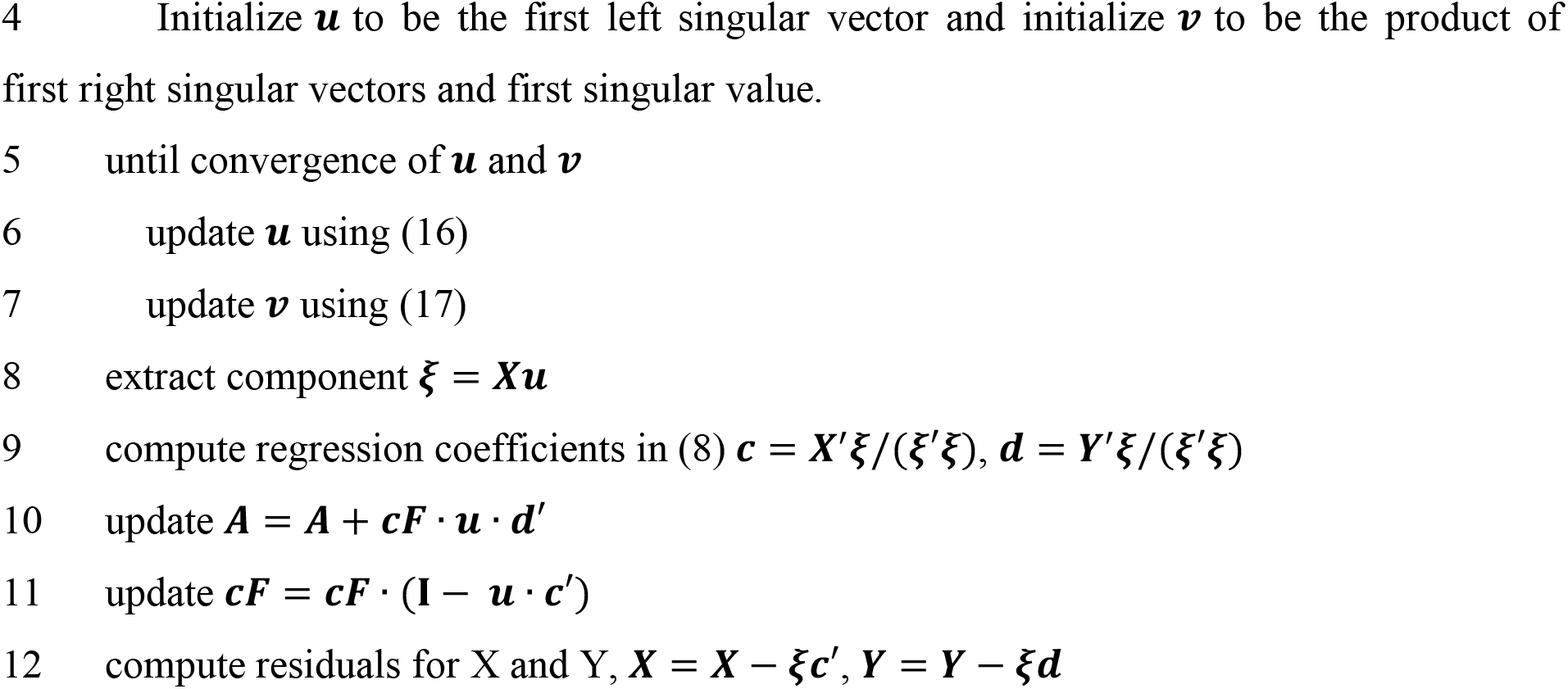
Huber-Berhu PLS regression

### Tuning criteria and choice of the PLS dimension

The Huber-Berhu PLS has two tuning parameters, namely, the penalization parameter *λ* and the number of hidden components *K*. To select the best penalization parameter, *λ*, a common k-fold cross-validation (CV) procedure that minimizes the overall prediction error is applied using a grid of possible values. If the sample size is too small, CV can be replaced by leave-one-out validation; this procedure is also used in [36, 49] for tuning penalization parameters.

To choose the dimension of PLS, the 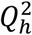 criteria was adopted. 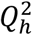 criteria were first proposed by Tenenhaus [50]; These criteria characterize the predictive power of the PLS model by performing cross-validation computation. 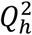 is defined as:

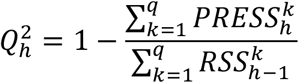

where 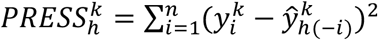 is the Prediction Error Sum of Squares, and 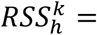 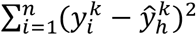 is the Residual Sum of Squares for the variable *k* and the PLS dimension *h*. The criterion for determining if *ξ*_*h*_ contributes significantly to the prediction is:

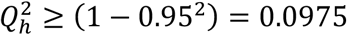

This criterion is also used in SIMCA-P software [51] and sparse PLS [37]. However, the choice of the PLS dimension still remains an open question. Empirically, there is little biological meaning when *h* is large and good performance appears in 2-5 dimensions.

## Results

### Validation of Huber-Berhu PLS with lignin biosynthesis pathway genes and regulators

The HB-PLS algorithm was examined for its accuracy in identifying lignin pathway regulators using the *A. thaliana* microarray compendium data set produced from stem tissues [39]. TFs identified by HB-PLS were compared to those identified by SPLS. The 50 most relevant TFs in the lignin biosynthesis pathway were identified using HB-PLS (Figure 3A) and compared to those identified by SPLS (Figure 3B), respectively. The positive lignin biosynthesis pathway regulators, which are supported by literature evidence, are shown in coral color. The HB-PLS algorithm identified 15 known lignin pathway regulators. Of these, MYB63, SND3, MYB46, MYB85, LBD15, SND1, SND2, MYB103, MYB58, MYB43, NST2, GATA12, VND4, NST1, MYB52, are transcriptional activators of lignin biosynthesis in the SND1-mediated transcriptional regulatory network [52], and LBD15 [53] and GATA12 [54] are also involved in regulating various aspects of secondary cell wall synthesis. Interestingly, SPLS identified the same set of pathway regulators as HB-PLS, though their ranking orders derived from connectivities to psthway genes are different.

**Figure 3.**
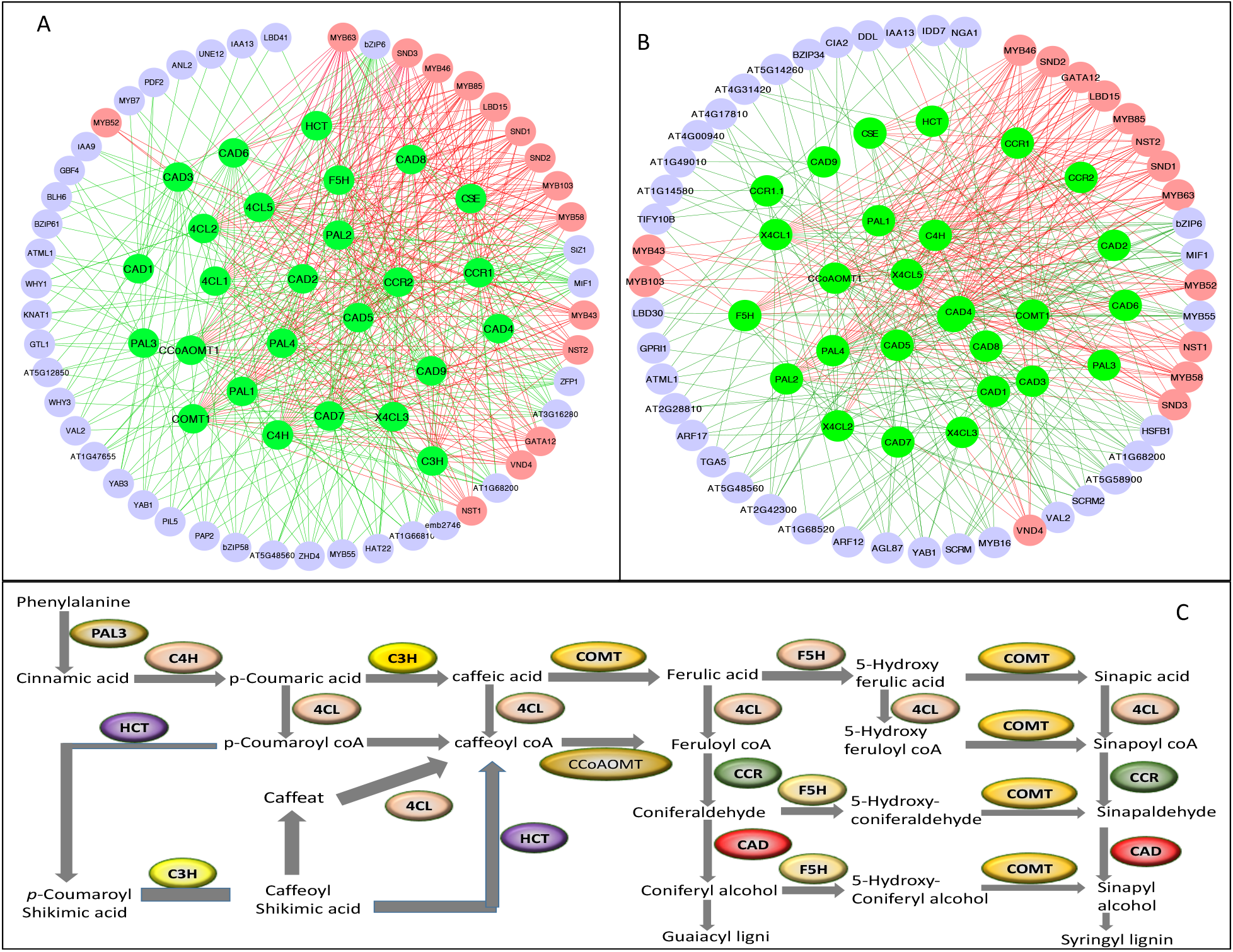
The 50 most important TFs in the lignin biosynthesis pathway (purple and coral nodes) were identified using Huber-Berhu-Partial Least Squares (HB-PLS) (A) and compared to those identified by sparse partial least squares (SPLS) (B). Green nodes (inside the circles) represent lignin biosynthesis genes. Coral nodes represent positive lignin pathway regulators identified in the literature, and shallow purple nodes contain other predicted transcription factors that do not have experimentally validated supporting evidence for the time being.

### The performance of HB-PLS with SPLS

Since lignin pathway regulators have been well characterized experimentally [55], they are specifically suited for determining the efficiency of the HB-PLS method. To do this, we selected two methods, SPLS and PLS, as comparisons. For each output TF list to a pathway gene yielded from one of three methods, we applied a series of cutoffs, with the number of TFs retained varying from 1 to 40 in a shifting step of 1 at a time, and then counted the number of positive regulatory genes in each of the retained lists. The results are shown in Figure 4.

**Figure 4.**
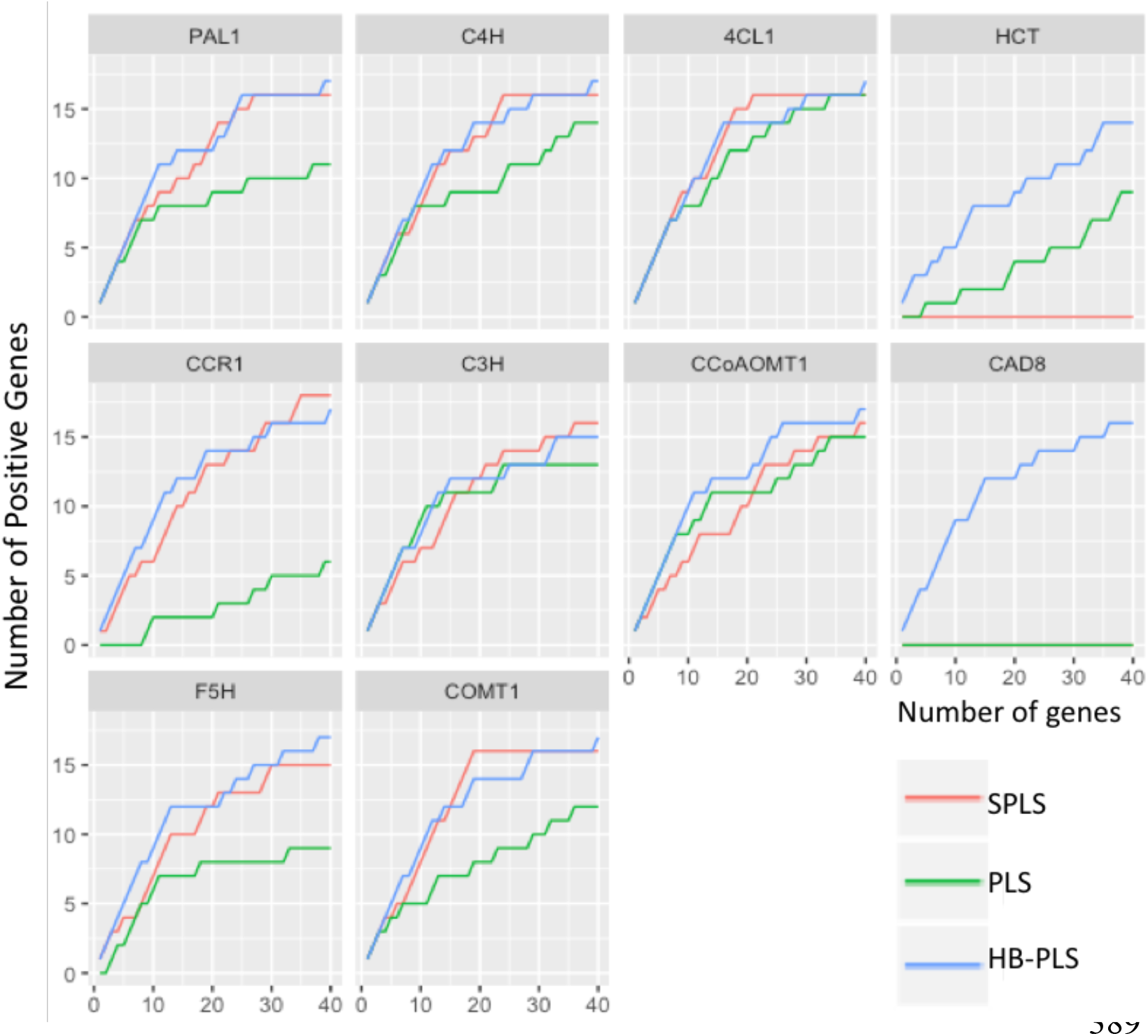
The performance of Huber-Berhu-Partial Least Squares (HB-PLS) was compared with the conventional partial least squares (PLS) and the sparse partial least squares (SPLS) method.

The results indicate that the HB-PLS and SPLS methods, in many cases, are much more efficient in recognizing positive regulators to a pathway gene compared to PLS method. For most pathway genes, like PAL1, C4H, CCR1, C3H, and COMT1, HB-PLS method could identify more positive regulators when the top cut-off lists contained fewer than 20 regulators compared to SPLS method. For HCT, CCoAOMT1, CAD8 and F5H, HB-PLS was almost always more efficient than SPLS when the top cut-off lists contained fewer than 40 regulators. For pathway gene CAD8, SPLS and PLS both failed to identify positive regulators while HB-PLS performed efficiently.

### Prediction of photosynthetic pathway regulators using Huber-Berhu PLS

Photosynthesis is mediated by the coordinated action of about 3,000 different proteins, commonly referred to as photosynthesis proteins [56]. In this study, we used genes from the photosynthesis light reaction (PLR) pathway to study which regulatory genes can potentially control photosynthesis. Analysis was performed using HB-PLS, with SPLS as a comparative method. The compendium data set we used is comprised of 63 and 36 RNA-seq data sets from maize leaves and seedlings, respectively. Expression data for 2616 TFs and 30 PLR pathway genes were extracted from the above compendium data set and used for analyses. The results of HB-PLS and SPLS methods are shown in Figure 5A and 5B, respectively. HB-PLS identified 14 positive TFs while SPLS identified 13 positive TFs. Among the 14 positive TFs identified by the HB-PLS method, NF-YC4 mediates light-controlled hypocotyl elongation by modulating histone acetylation [57]. The chloroplast psbD gene encodes the D2 protein of the photosystem II (PSII) reaction center. In the green alga *Chlamydomonas reinhardtii*, D2 synthesis requires a high-molecular-weight complex containing the RNA stabilization factor NAC2 [58]. GBF6 is indicated to control CLPB3 in an irradiance-dependent manner [59]; CLPB3 encodes a molecular chaperone involved in plastid differentiation mediating internal thylakoid membrane formation [60]. The chloroplast protein phosphatase TAP38/PPH1 is required for efficient dephosphorylation of the light-harvesting complex II (LHCII) anthenna and the state transition from state 2 to state 1 [61]. The transcription factor bZIP63 is required for adjustment of circadian period by sugars [62]. PIF1 negatively regulates chlorophyll biosynthesis and seed germination in the dark, and light-induced degradation of PIF1 relieves this negative regulation to promote photomorphogenesis [63]. The transcription factor HY5 is a key regulator of light signaling, acting downstream of photoreceptors. HY5 also binds sites in the promoter of the STOMAGEN (STOM) gene, which encodes a peptide regulator of stomatal development [64]. HY5 also binds and regulates the circadian clock gene PRR7, which affects the operating efficiency of PSII under blue light [65]. By QTL mapping, WRKY2 and PRR2 are predicted regulators that control photosynthesis [66]. mtTTF is induced by light (particularly blue light) [67]. mTERF6 is required for photoautotrophic growth early in development, and mterf6-5 exhibited reduced growth and defective chloroplasts [68]. Of the TFs identified by SPLS, mTTF-2 is induced by light (particularly blue light) [67]. REB3 and WRKY11 are predicted TFs that control photosynthesis through QTL mapping [66].

**Figure 5.**
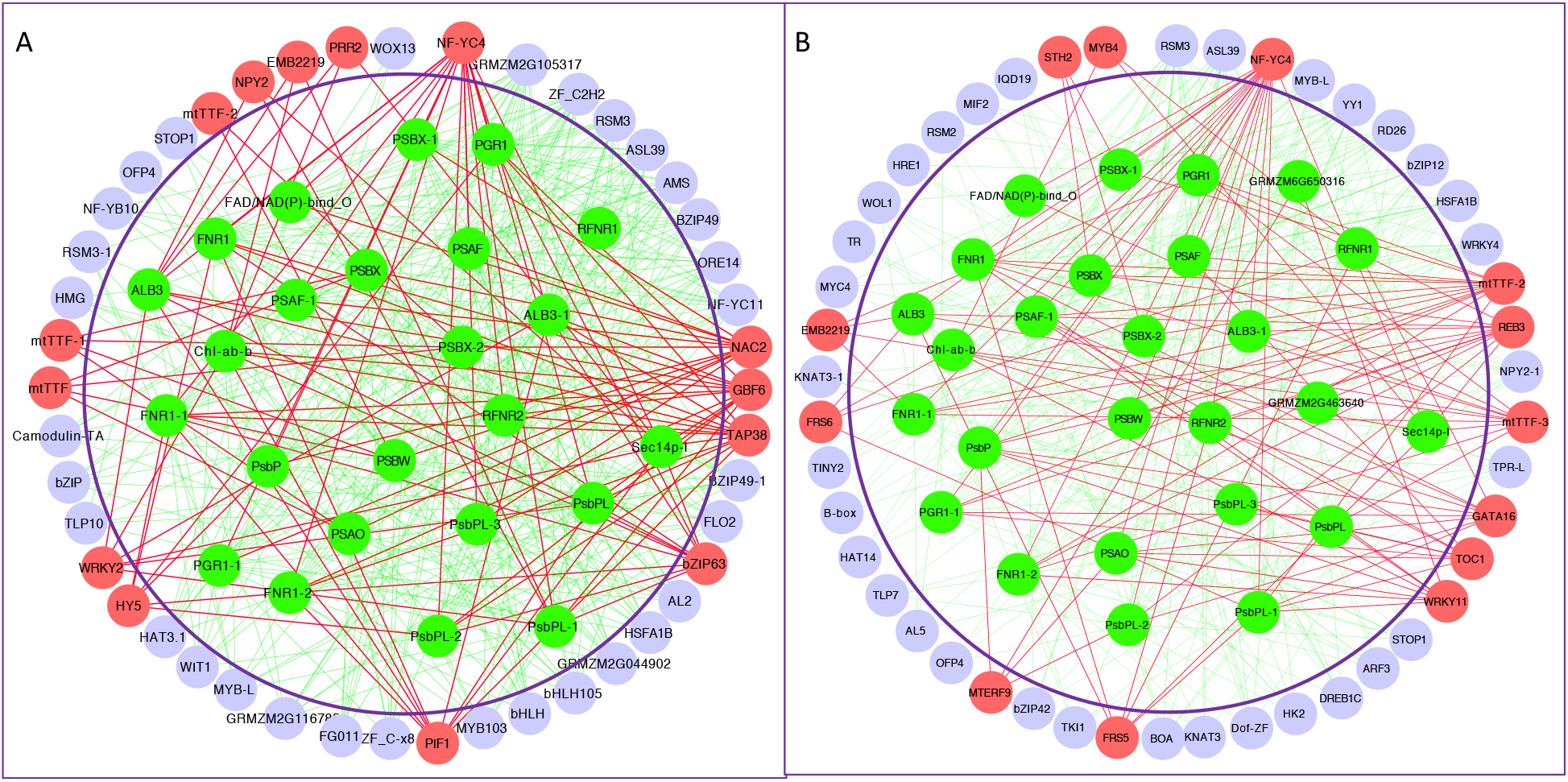
The performance of Huber-Berhu-Partial Least Squares (HB-PLS) (A) was compared with the sparse partial least squares (SPLS) method (B) in identifying regulators that affects maize photosynthesis light reaction (PLR) pathway genes. The green nodes represent photosynthesis light reaction pathway genes. Coral nodes represent predicted biological process or pathway regulators, and shallow purple nodes contain other predicted TFs that do not have experimentally validated supporting evidence for the time being.

GATA16 controls greening, hypocotyl elongation [69]; TOC1 mis-expressing plants were shown to have altered ABA-dependent stomata closure [70]. FRS5 is expressed in cotyledons of light-grown seedlings, which is consistent with a potential role for FRS5 in regulating light control in *Arabidopsis* development [71]. Moreover, FRS6 likely acts as a positive regulator in the phyB signaling pathway controlling flowering time [71]. MTERF9 has a significant role in light signaling as well as in aminoacylation and seed storage [71]. EMB2219 encodes a mitochondrial transcription termination factor that is localized and enriched in proplastids and chloroplasts [72]. STN2 enhances the rate of photosynthesis and alleviates photoinhibition in *Solanum tuberosum* [73]. Vannini et al. (2004) have reported that *Arabidopsis* plants overexpressing OsMYB4 show improved PSII stability and higher tolerance to photoinhibition [74].

## Discussion

The identification of gene regulatory relationships by the construction of gene regulatory networks from gene expression data sets has inherent challenges due to high dimensionality and multicollinearity. High dimensionality is caused by a multitude of gene variables while multicollinearity is largely the result of a large number of genes in a relatively small number of samples. One method that can circumvent these challenges is partial least squares (PLS), which couples dimension reduction with a regression model. However, because PLS is not particularly suited for variable/feature selection, it often produces linear combinations of the original predictors that are hard to interpret due to high dimensionality [75]. To solve this problem, Chun and Keles developed an efficient implementation of sparse PLS, referred to as the SPLS method, based on the least angle regression [76]. SPLS was then benchmarked by means of comparisons to well-known variable selection and dimension reduction approaches via simulation experiments [75]. We used the SPLS method in our previous study [41] and found that it was highly efficient in identifying pathway regulators and thus can be used as a benchmark for the development of new algorithms.

In this study, we developed a PLS regression that incorporates the Huber loss function and the Berhu penalty for identification of pathway regulators using gene expression data. Although the Huber loss function and the Berhu penalty have been proposed in regularized regression models [43, 77], this is the first time that both have been used in the PLS regression at the same time. The Huber function is a combination of linear and quadratic loss functions. In comparison with other loss functions (e.g., square loss and least absolute deviation loss), Huber loss is more robust to outliers and has higher statistical efficiency than the LAD loss function in the absence of outliers. The Berhu function [33] is a hybrid of the *ℓ*_2_ penalty and the *ℓ*_1_ penalty. It gives a quadratic penalty to large coefficients and a linear penalty to small coefficients. Therefore, the Berhu penalty has advantages of both the *ℓ*_2_ and *ℓ*_1_ penalties: smaller coefficients will tend to shrink to zero while the coefficients of a group of highly correlated predictive variables will not be changed much if their coefficients are large.

A comparison of HB-PLS with SPLS suggests that they have comparable efficiencies. The implementation of HB-PLS and SPLS algorithms for identifying lignin pathway regulators in *Arabidopsis* led to the identification of 15 positive regulators using each algorithm, and implementation of the HB-PLS and SPLS algorithms for identifying PLR pathway regulators in maize resulted in 14 and 13 positive regulators, respectively. The HB-PLS and SPLS algorithms each performed better than the conventional PLS method in identifying positive pathway regulators. The simulation of performance efficiency of both methods for each of the lignin pathway genes suggests that HB-PLS identifies more positive regulators in the top of output lists of pathway regulatros that have fewer than 20 TFs. However, as output regulatory gene lists increase to more than 20 genes, so does the efficiency of SPLS. In the output lists of HB-PLS and SPLS, the positive regulators share some common genes but their rankings are different, indicating that the two algorithms have unique specificities that can be used to identify different sets of positive pathway regulators through modeling GRNs.

## Conclusions

A proximal gradient descent algorithm was developed to solve a regression optimization problem. In this regression, the Huber function was used as the loss function and the Berhu function was used as the penalty function. An optimal one-dimensional clustering algorithm was adopted to cluster the regression coefficients and then the elbow point was used to determine the non-zero variables. The Huber function is more robust in dealing with outlier and non-Gaussian error while the Berhu function integrates the advantages of both *ℓ*_1_ and *ℓ*_2_ penalties. The group effect of the Huber-Berhu regression makes it suitable for modeling transcriptional regulatory relationships. Simulation results showed that the Huber-Berhu regression has better performance in identifying non-zero variables. When modeling the regulatory relationships from TFs to a pathway, HB-PLS is capable of dealing with the high multicollinearity of both TFs and pathway genes. Implementation of the HB-PLS to *Arabidopsis* and maize data showed that HB-PLS can identify comparable numbers of positive TFs in the two pathways tested. However, there were differences in the pathway regulators identified and their rankings; in particular, positive TFs tended to be present in highly ranked positions in output lists. This is an advantage for selecting candidate regulators for experimental validation. Our results indicate that HB-PLS will be instrumental for identifying novel biological process or pathway regulators from high dimensional gene expression data.

## Contributions

WD developed the methods and implemented the method in R. HW, SL, KZ are involved in desiging and improving the method. CH, ZW and LW were involved in data collection and network construction, interpretation, and plotting. WD, HW and SL wrote the manuscript. KZ, ZW, SL and HW revised the manuscript.

## Acknowledgements

NSF Plant Genome Program [1703007 to SL and HW]; NSF Advances in Biological Informatics [dbi-1458130 to HW]; USDA McIntire-Stennis Fund to HW.

